# The HIV-1 capsid serves as a nanoscale reaction vessel for reverse transcription

**DOI:** 10.1101/2023.11.08.566350

**Authors:** Jordan Jennings, Harrison Bracey, Danny T. Nguyen, Rishav Dasgupta, Alondra Vázquez Rivera, Nicolas Sluis-Cremer, Jiong Shi, Christopher Aiken

## Abstract

The viral capsid performs critical functions during HIV-1 infection and is a validated target for antiviral therapy. Previous studies have established that the proper structure and stability of the capsid are required for efficient HIV-1 reverse transcription in target cells. Moreover, it has recently been demonstrated that permeabilized virions and purified HIV-1 cores undergo efficient reverse transcription in vitro when the capsid is stabilized by addition of the host cell metabolite inositol hexakisphosphate (IP6). However, the molecular mechanism by which the capsid promotes reverse transcription is undefined. Here we show that wild type HIV-1 particles can undergo efficient reverse transcription *in vitro* in the absence of a membrane-permeabilizing agent. This activity, originally termed “natural endogenous reverse transcription” (NERT), depends on expression of the viral envelope glycoprotein during virus assembly and its incorporation into virions. Truncation of the gp41 cytoplasmic tail markedly reduced NERT activity, indicating that gp41 permits the entry of nucleotides into virions. Protease treatment of virions markedly reduced NERT suggesting the presence of a proteinaceous membrane channel. By contrast to reverse transcription in permeabilized virions, NERT required neither the addition of IP6 nor a mature capsid, indicating that an intact viral membrane can substitute for the function of the viral capsid during reverse transcription *in vitro*. Collectively, these results demonstrate that the viral capsid functions as a nanoscale container for reverse transcription during HIV-1 infection.

## Introduction

The HIV-1 capsid is a conical shell composed of a single viral protein (CA) assembled as a lattice of approximately 1200 subunits (reviewed in (1)). The capsid surrounds the ribonucleoprotein complex (RNP) which consists of the viral genomic RNA, tRNA^Lys^, nucleocapsid protein (NC), reverse transcriptase (RT) and integrase (IN) enzymes. The capsid plays multiple roles during infection and is an emerging target for therapy (2, 3). The structure and stability of the capsid are critical for HIV-1 reverse transcription in target cells. Mutations that destabilize the capsid or perturb its structure result in impaired reverse transcription (4–10), as does targeting by restrictive variants of the host protein TRIM5α (reviewed in (11)). By contrast, hyperstabilization of the capsid via mutations or antiviral compounds can inhibit nuclear entry without impairing reverse transcription, as can amino acid substitutions in CA that disrupt binding of host proteins to the capsid (12–14). Antiviral compounds targeting CA can also inhibit reverse transcription, nuclear entry, and integration, reflecting involvement of the capsid in these three early steps of infection (15–18).

The importance of capsid stability has also been underscored by studies of reverse transcription in purified HIV-1 cores or permeabilized virions *in vitro*. In the absence of a capsid-stabilizing agent, such endogenous reverse transcription (ERT) reactions with HIV-1 are inefficient, resulting in synthesis of full-length viral DNA in only a small fraction of viral cores. Addition of the host cell metabolite IP6 dramatically enhances the synthesis of late reverse transcription products (19, 20). ERT activity is correlated with capsid stabilization by IP6, further emphasizing the role of a stable capsid in reverse transcription (20). Addition of capsid-destabilizing antiviral compounds inhibited the reaction, but this was partially overcome by increasing the concentration of IP6 (21). Although depletion of both IP6 and inositol pentakisphosphate (IP5) showed minimal effects on HIV-1 infection of target cells (22), it potentiated inhibition of infection by PF74 and the highly potent antiviral drug Lenacapavir (23), suggesting that target cell inositol phosphates help stabilize the capsid following entry of the viral core into target cells.

Despite the well-established link between HIV-1 capsid integrity and reverse transcription, the mechanism by which the capsid promotes completion of reverse transcription is unknown. Reverse transcriptase is a low processivity enzyme that dissociates after synthesis of short stretches of DNA (reviewed in (24)). Thus, a plausible hypothesis is that the capsid serves as a container for reactants and catalysts, ensuring that a sufficient local concentration of RT is maintained during the reaction (Fig 1A). Additionally, the capsid could function as a molecular scaffold that serves to concentrate the reactants in a manner akin to surface catalysis of chemical reactions (Fig. 1B). While these two models are not mutually exclusive, the container model implies that the capsid must be sealed during the reaction, whereas a partially intact capsid could function as a scaffold. Here we report evidence resulting from the unexpected observation that intact HIV-1 particles can undergo efficient reverse transcription. We show that, by contrast to permeabilized virions or purified HIV-1 cores, reverse transcription in nonpermeabilized virions occurs in the absence of a mature viral capsid, is insensitive to capsid-destabilizing mutations and compounds, and does not require the addition of IP6.

**Fig 1.**
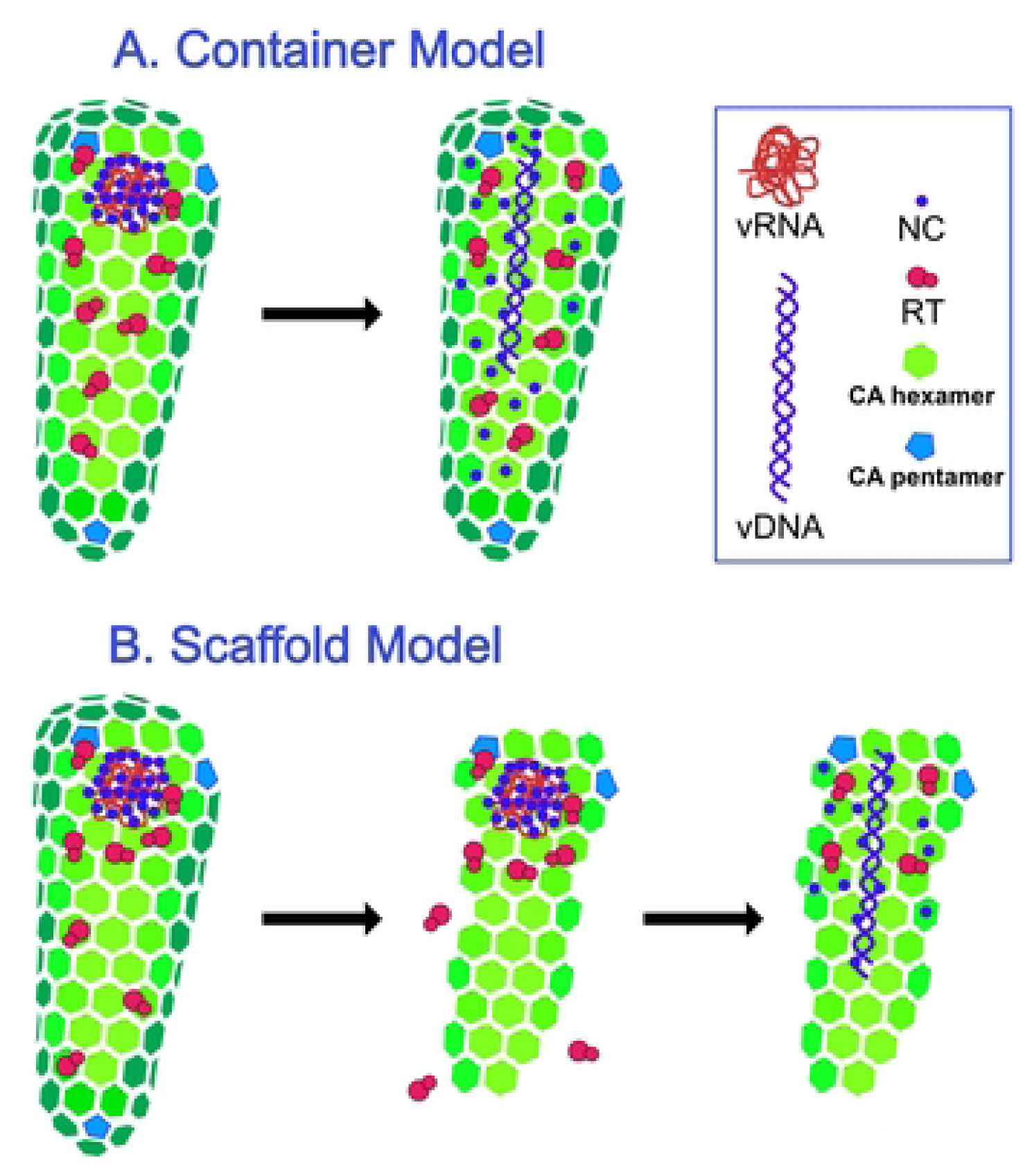
Hypothetical models for the role of the HIV-1 capsid in reverse transcription. A. The container model posits that the capsid serves to maintain the concentrations of proteins and nucleic acids required for reverse transcription and predicts that the capsid must remain fully intact during the reaction. B. In the scaffold model, the inner face of the assembled capsid functions as a platform on which the reaction takes place, suggesting that the capsid may be semi-intact.

## Results

### Capsid stabilization by IP6 results in DNA synthesis that is resistant to added nuclease

Previous studies have indicated that stabilization of the viral capsid by nucleotides and/or IP6 promotes ERT activity *in vitro* (19, 20, 25, 26). In principle, the viral capsid could serve as a container for the reaction, a scaffold on which the reaction occurs, or both. The container hypothesis posits that the capsid is sealed and therefore should exclude molecules that exceed the size of the pores in the capsid lattice (i.e., average-sized proteins). To test this, we performed ERT reactions in the presence and absence of DNase I and various concentrations of IP6 to result in graded levels of capsid stabilization. In the absence of DNase I, ERT was progressively enhanced by concentrations of IP6 of up to 10 μM (Fig 2, reactions 1-5). As previously reported (19, 20), the greatest increase was observed with synthesis of the late reverse transcripts. In parallel reactions containing DNase I, minimal effects were observed in reactions containing 1 and 10 uM IP6, the concentration range in which the reaction was most efficient (Fig 2, reactions 6-9). As a control to determine whether capsid disassembly results in susceptibility of nascent DNA to degradation, we included reactions in which DNase I and the capsid-destabilizing compound PF74 were added after 4h and incubated for an additional 1h (Fig 2, reactions 10-12). In these reactions, the level of ERT was markedly reduced, indicating that capsid destabilization by PF74 rendered the reverse transcripts sensitive to degradation. These results indicated that IP6 stabilization of the capsid is associated with protection of the reverse transcribed DNA from nuclease degradation, suggesting that the capsid can sequester the DNA products. However, they did not exclude a possible scaffolding function for the capsid.

**Fig 2.**
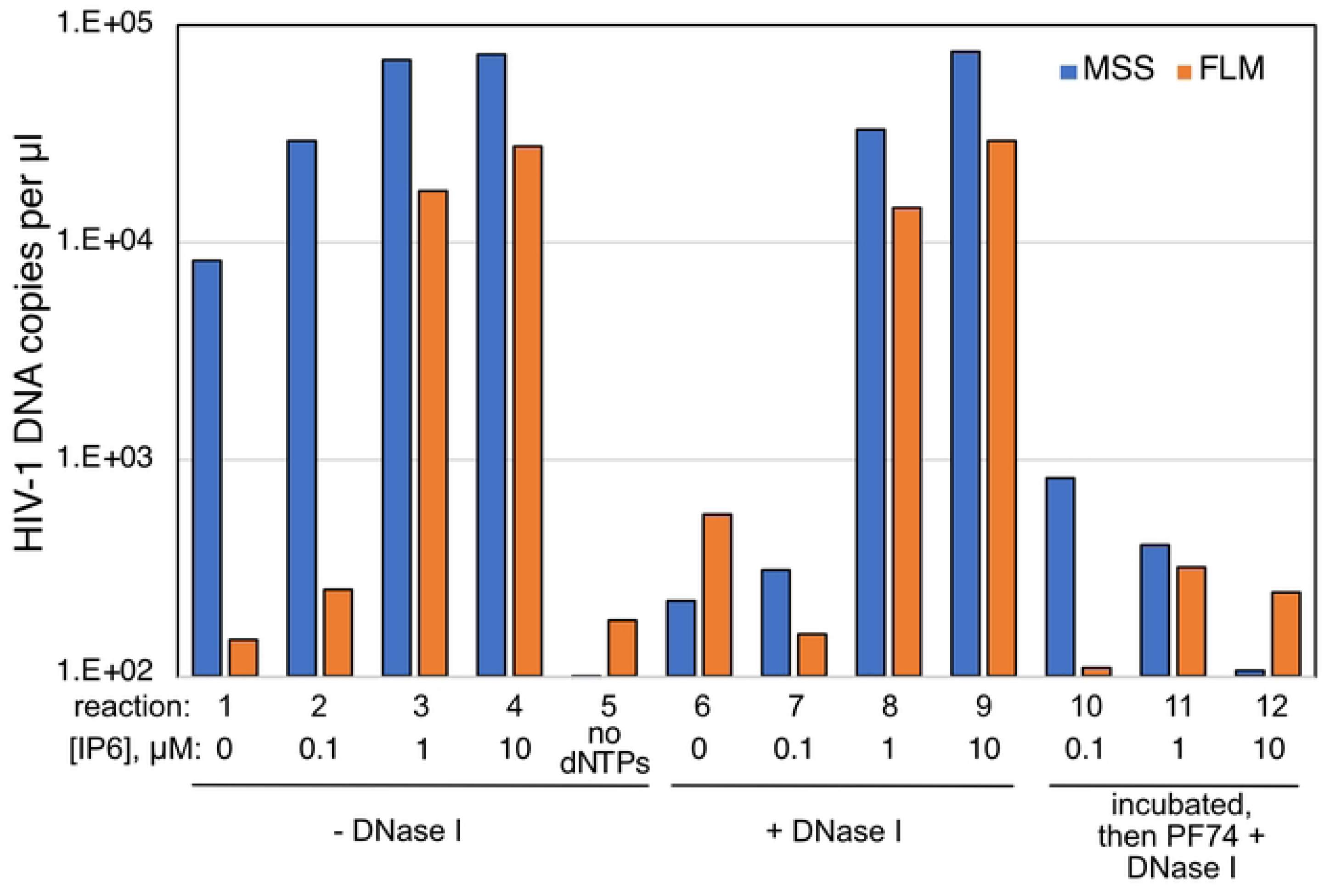
IP6-dependent ERT reactions are resistant to degradation by DNase I. ERT reactions were performed with HIV-1 particles in the presence of the indicated concentrations of IP6, with or without added DNase I. Reactions were incubated for 4h. In reactions 10-12, PF74 was added to a concentration of 20 μM and DNAse I to 20 μg/ml, and the reactions were incubated for an additional 60 min. Products were purified and quantified by qPCR using primers specific for minus strand strong stop (MSS) and full length minus strand (FLM) amplicons. Results shown are representative of two independent experiments.

### Efficient reverse transcription can occur in nonpermeabilized HIV-1 particles

In the course of this work, we unexpectedly observed that HIV-1 reverse transcription occurred in reactions lacking detergent (Triton X-100) and IP6 and was nearly as efficient as in reactions containing both components (Fig 3A). High quantities of both early (minus strand strong stop; MSS) and late (full length minus; FLM) products were produced in the reactions lacking detergent. As in reactions containing detergent and IP6, ERT reactions lacking detergent were inhibited by nucleoside and non-nucleoside reverse transcription inhibitors and by aldrithiol-2 (AT-2), a compound that inactivates the nucleocapsid protein by oxidizing zinc-coordinating Cys residues (27) (Fig 3B). However, by contrast to reactions containing detergent or purified cores, the reaction with nonpermeabilized virions was not inhibited by PF74. These observations suggested that wild type HIV-1 particles are naturally permeable to dNTPs and that the nonpermeabilized ERT reaction lacks the capsid stability requirement observed in reactions involving either permeabilized HIV-1 virions or purified cores.

**Fig 3.**
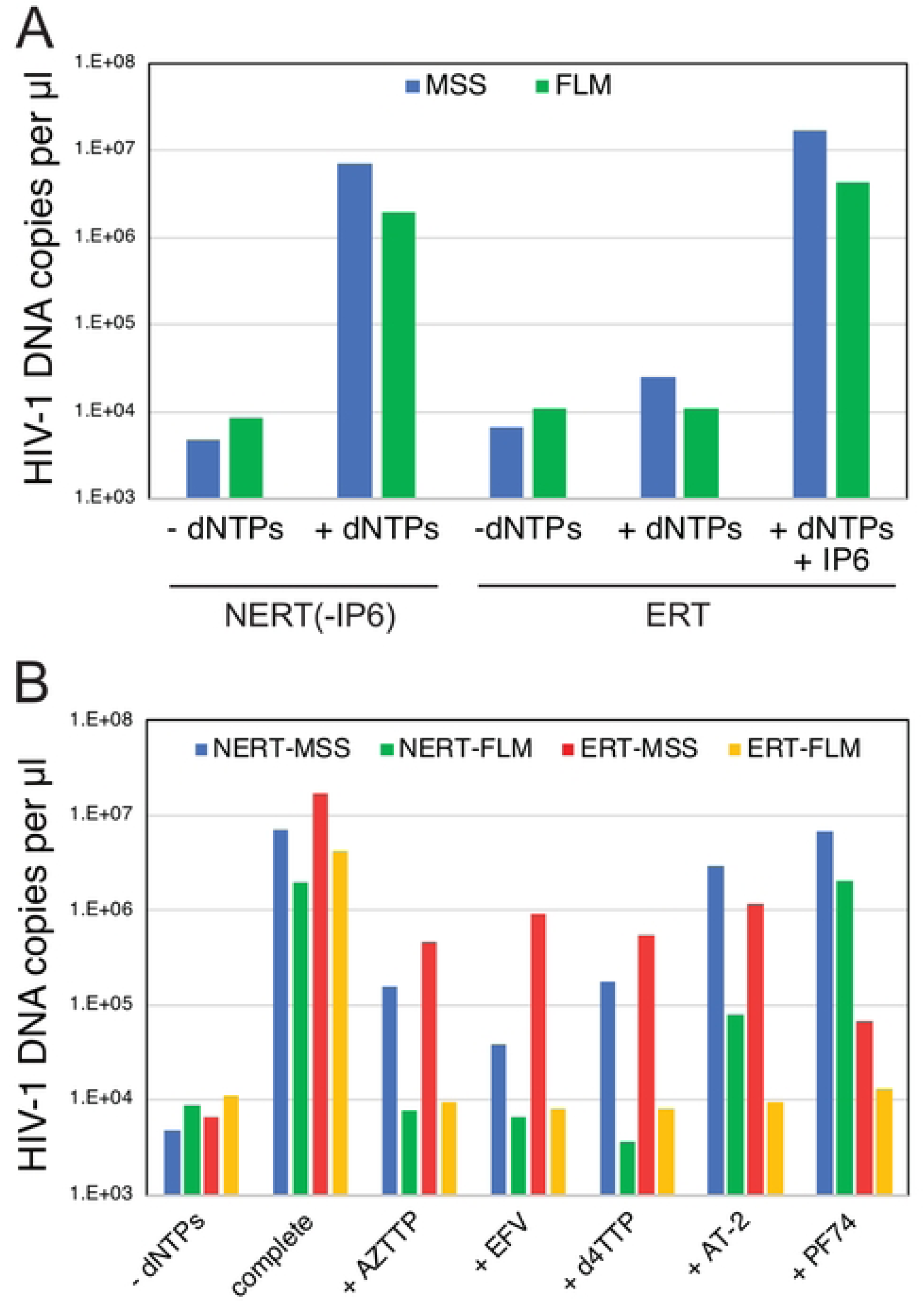
Reverse transcription in nonpermeabilized HIV-1 particles does not require addition of IP6 and is resistant to PF74. Reactions were incubated for 14h. A. Reactions with wild type HIV-1 particles were performed in the presence or absence of dNTPs (dATP, dCTP, dGTP, and TTP; 0.1 mM each), 0.1% Triton X-100, and 0.1 mM IP6. B. Reactions were performed in the absence or presence of detergent and IP6 (NERT and ERT, respectively) with the indicated inhibitors: 10 μM azidothymidine triphosphate (AZTTP); 1 μM efavirenz (EFV); 10 μM stavudine triphosphate (d4TTP); 1 mM aldrithiol (AT-2); 10 μM PF-3450074 (PF74). The reaction designated “complete” contained dNTPs but no inhibitor. Results shown are representative of two independent experiments.

### NERT is dependent on the viral Env glycoprotein complex

An earlier study linked the ability of nonpermeabilized virions to undergo reverse transcription *in vitro* to the viral Env protein gp41 (28). This activity was termed Natural Endogenous Reverse Transcription (NERT). To confirm the previous findings, we performed NERT and ERT reactions containing wild type and Env-deficient particles and a series of Env truncation mutants lacking portions of the gp41 cytoplasmic tail (CT). As previously reported, Env-deficient particles were markedly impaired in NERT activity relative to wild type virions, with an approximately 50-fold reduction in synthesis of late products (Fig 4A). By contrast, the lack of Env had no effect on ERT. Truncation of 28 or 42 amino acids (CT28 and CT42, respectively) from the carboxyl terminus of gp41 only moderately reduce NERT activity relative to the level observed in wild type particles. However, removal of the C-terminal 93 amino acids (CT93) markedly reduced NERT, as did the more extensive truncations of 104 and 144 amino acids (CT104, CT144). Immunoblot analysis of pelleted virions showed that the five truncated Env proteins were incorporated into HIV-1 particles and that the mutants that exhibited impaired NERT activity contained at least as much gp41 protein as wild type particles (Fig 4, panels B and C). These results confirmed the earlier report that the cytoplasmic tail of gp41 plays a role in permeabilizing HIV-1 particles to dNTPs (28); further, they define a central region of the cytoplasmic tail as necessary for this activity.

**Fig 4.**
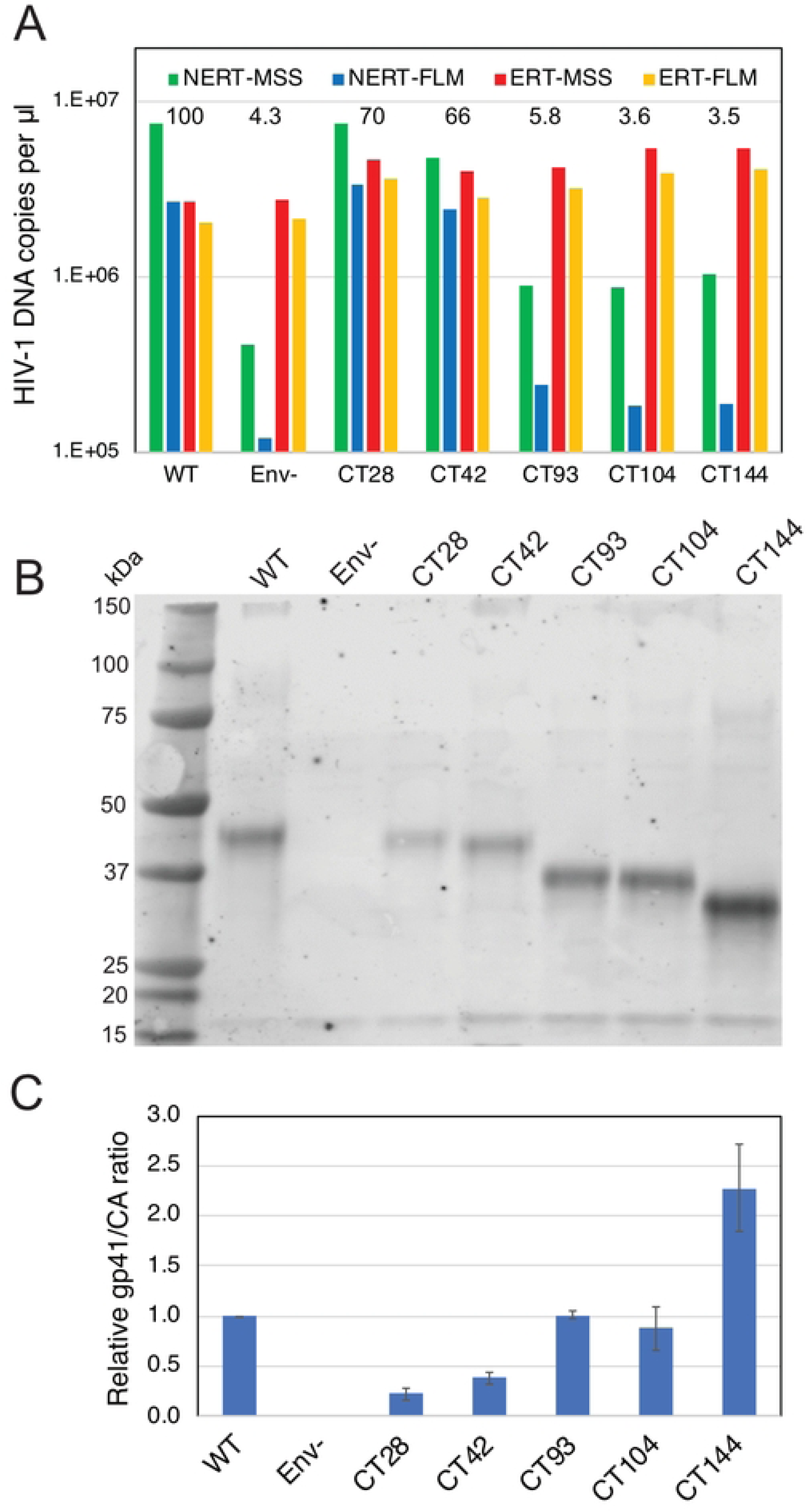
Reverse transcription in nonpermeabilized HIV-1 particles requires the gp41 CT. A. Reactions containing wild type (WT), Env-deficient (Env^-^), and the indicated gp41 C-terminal truncation mutants were performed in the absence (NERT) and presence (ERT) of detergent and IP6. Reactions were incubated for 14h and the early (MSS) and late (FLM) products quantified by qPCR. Numerical values in the graph represent the NERT to ERT ratio of FLM product levels in each reaction relative to that measured for wild type virions. B. Pelleted virions used in the reactions shown in A were analyzed by immunoblotting using a monoclonal antibody recognizing a membrane-proximal epitope in the gp41 CT. C. Ratio of band intensities of gp41 and CA shown in B. Error bars represent the range of values from two independent experiments. Results shown in each panel are representative of two independent experiments.

We also asked whether Env proteins from other viruses can support NERT. Pseudotyped HIV-1 particles bearing the Env proteins of amphotropic murine leukemia virus (A-MLV) exhibited NERT activity approximately 22% that of particles bearing HIV-1 Env (Fig 5A). By contrast, HIV-1 particles bearing the vesicular stomatitis virus glycoprotein (VSV-G) exhibited only 5% of the activity of HIV-1 Env-bearing particles. Assays of particles released from 293T cells transfected with a full-length molecular clone of SIVmac293 also revealed that the virions were also active in NERT reactions and that the Env protein was required (Fig 5B). These results demonstrate that the A-MLV and SIVmac239 Env glycoproteins, but not VSV-G, permit dNTP entry into nonpermeabilized virions, albeit with varying efficiencies. NERT activity was also exhibited by particles produced from several cloned HIV-1 primary isolate Env proteins, with efficiencies ranging from 27% to 119% of their corresponding ERT values (Fig 5C).

**Fig 5.**
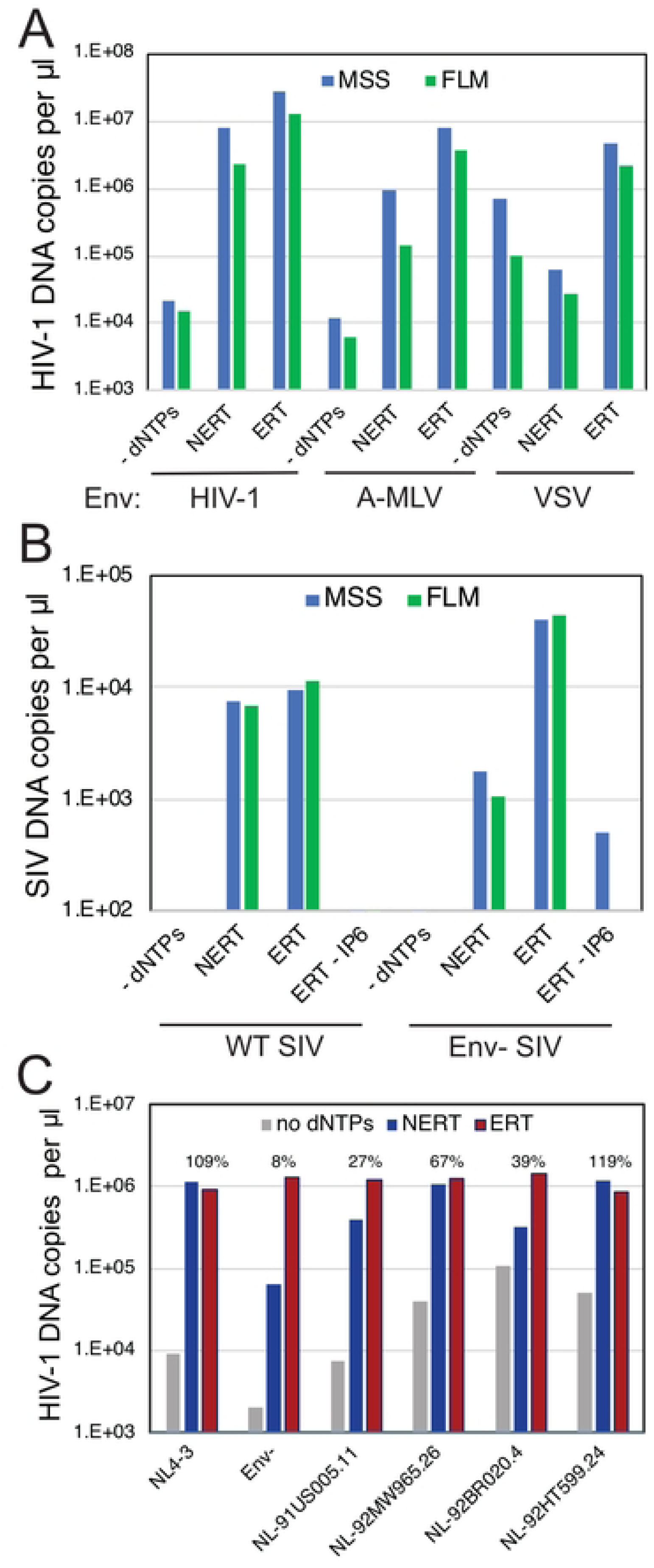
NERT activity depends on the viral Env protein. NERT and ERT reactions were performed with the indicated viruses. A. Analysis of wild type HIV-1 particles (HIV-1), A-MLV pseudotyped HIV-1 particles (A-MLV), and VSV-G-pseudotyped HIV-1 particles (VSV). B. Analysis of wild type and Env-SIV_mac_239 particles. C. Analysis of wild type (NL4-3), Env-, and NL4-3 chimerae encoding Env proteins from HIV-1 primary isolates. Shown are the levels of late product synthesis detected in each NERT and ERT reaction. Results are representative of two independent experiments.

### NERT activity is exhibited by viruses produced in T cells

To determine whether NERT activity is limited to HIV-1 particles released from transfected 293T cells, we assayed NERT in wild type and Env-virions harvested from infected T cells. Five T cell lines (CEM, H9, Jurkat, MT-4, and SupT1) were inoculated with wild type and Env-particles that also bore the VSV-G protein to enhance initial infection. Cells were washed and cultured for 4 days, after which the supernatants were collected, treated with DNase I to remove potential contaminating plasmid DNA, and concentrated by pelleting through 20% sucrose to remove the nuclease. The resulting stocks were assayed in parallel for NERT and ERT activity. We observed high levels of NERT activity in the wild type virions released from CEM, Jurkat, and SupT1 cells (Fig 6). Wild type particles released from H9 and MT-4 cells exhibited lower levels of NERT. Env-defective particles released from all five cell types exhibited minimal NERT relative to ERT activity. These results indicated that HIV-1 particles released from some human T cell lines are competent for NERT and that the activity is dependent on the Env protein.

**Fig 6.**
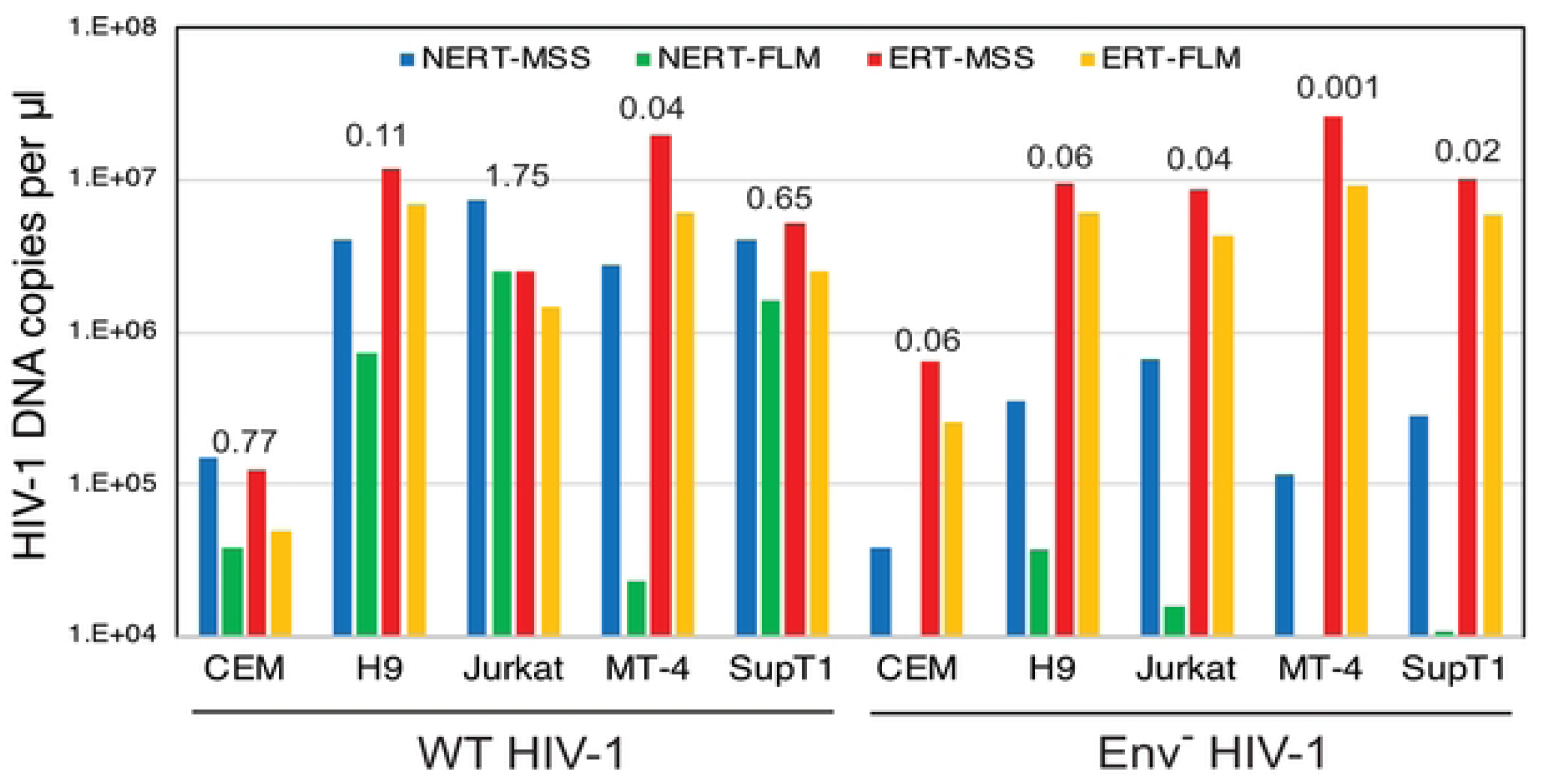
NERT efficiency in HIV-1 particles varies with the cellular source of the virions. Wild type and Env-HIV-1 particles were harvested from the indicated T cell lines, concentrated, and assayed for NERT and ERT activity. Shown are the levels of early (MSS) and late (FLM) products. The numerical values above each sample represent the ratio of late product levels in the corresponding NERT and ERT reactions. Results shown are representative of two independent experiments.

### Minimally processed virions also exhibit NERT activity

Our standard procedure for producing virions involved transfection of cells, clarification of culture supernatants by low-speed centrifugation, filtration through a 0.45 μm pore-size syringe filter to remove organelles and large vesicles, digestion of remaining plasmid DNA by treatment with DNase I, pelleting by ultracentrifugation to concentrate the particles, and resuspension of the viral pellet by repeated pipetting. The extensive processing of the virions we employed left open the possibility that the dNTP membrane permeability induced by Env proteins could result from microdamage to the membrane induced during virion processing and storage.

To test whether physical damage of virions contributed to NERT activity, we performed reactions using virions that were minimally processed. Fresh culture supernatants were harvested from transfected cells and clarified by low-speed centrifugation without filtration. The virions were neither frozen, pelleted, nor otherwise concentrated. However, it was necessary to treat the virions with DNAse I to reduce the residual carryover plasmid DNA from the transfections. Nonetheless, the nuclease was not removed prior to the reactions to minimize the possibility of damaging the membrane upon pelleting of the virions. NERT reactions were performed in the presence and absence of dNTPs to distinguish nascent from residual plasmid DNA. After a 4h incubation, reactions were halted by the addition of EDTA, and the virions were then pelleted by ultracentrifugation at 4°C through a 20% sucrose cushion. DNA was purified and early and late products quantified by qPCR; values were normalized by the levels exogenous RT activity present in each virus stock. DNA synthesis in NERT reactions with these minimally processed virions was comparable to that observed with virions prepared by our normal procedure (Fig 7A). As expected, unprocessed Env-deficient particles exhibited substantially lower levels of NERT activity. These results show that minimally processed HIV-1 particles also support efficient NERT, suggesting that membrane damage during virion processing is an implausible explanation for efficient dNTP entry into virions.

**Fig 7.**
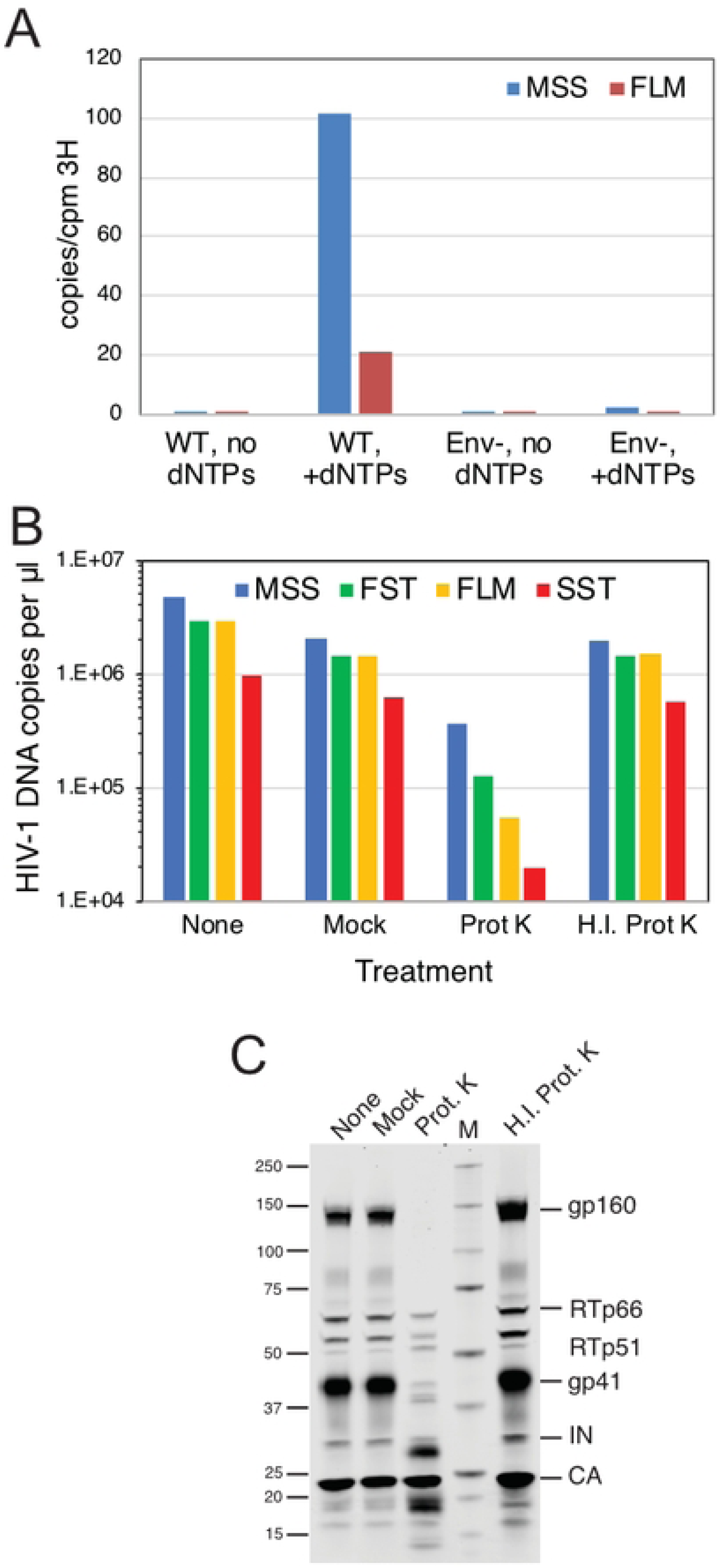
Treatment of intact HIV-1 particles with protease reduces NERT activity. A. NERT reactions in minimally processed wild type (WT) and Env-particles. Shown are the levels of early and late product synthesis in reactions containing and lacking dNTPs. B. NERT activity in concentrated virions after treatment for 1h with proteinase K (Prot K) or heat-inactivated proteinase K (H.I. Prot K). Reactions containing virions that were not incubated (None) and virions incubated with no proteinase K (Mock) served as controls. C. Immunoblot analysis of virions used in the reactions shown in B. The blot was probed with an antibody to the gp41 CT and with HIV-Ig, which recognizes RT, IN, and CA. Results shown are from one of two independent experiments.

### Protease treatment of HIV-1 particles reduces NERT activity

The ability of nonpermeabilized HIV-1 particles to undergo NERT suggested the presence of a membrane channel protein on HIV-1 particles that permits entry of dNTPs. To determine whether virion surface proteins render HIV-1 particles permeable to dNTPs, we assayed NERT in wild type virions following treatment with a nonspecific protease (proteinase K) to remove proteins from the virion surface. Treatment with active protease reduced late product synthesis in NERT reactions by more than 90% (Fig 7B). As a control for unexpected effects of protease treatment, we tested an identical concentration of proteinase K that had been heat inactivated. This treatment produced a negligible reduction of both ERT and NERT activity levels relative to the mock-treated control. Immunoblot analysis of the protease-treated particles showed a band profile reflecting cleavage of gp160 and gp41, with appearance of two faster-migrating bands detected with a monoclonal antibody against a membrane-proximal epitope in the gp41 CT (Fig 7C). We also noted a slight reduction in the intensity of bands corresponding to RT polypeptides following protease treatment of HIV-1 particles. By contrast, the level of pelleted CA protein was not affected. These results indicate that cleavage of proteins from the surface of the virus reduces ERT activity, supporting the hypothesis that a protein-dependent membrane channel is required for NERT activity.

In principle, the observed reduction in NERT activity upon protease treatment of virions could result from cleavage of either a viral or a cellular protein on the virion surface. To further probe the involvement of the gp41 CT in NERT activity, we examined two mutants encoding substitutions in the membrane-proximal region of the CT that result in protease cleavage of the tail (P203L and S205L; numbered according to the amino acid sequence of gp41). These viruses had been previously identified by selection for HIV-1 resistance to the cholesterol-binding compound amphotericin B methyl ester (AME) (29). Resistance to AME had been previously observed in CT truncation mutants (30), and acquisition of resistance via cleavage of the tail by the viral protease further supported the conclusion that AME sensitivity was conferred by the gp41 CT. We observed that NERT activity was markedly reduced in the P203L and S205L mutants, consistent with the requirement for the gp41 CT in NERT activity (Fig 8, panels A and B). Immunoblot analysis of the concentrated particles confirmed cleavage of a substantial portion of gp41 (Fig 8C). Thus, cleavage of the gp41 CT by the viral protease was associated with reduced NERT activity.

**Fig 8.**
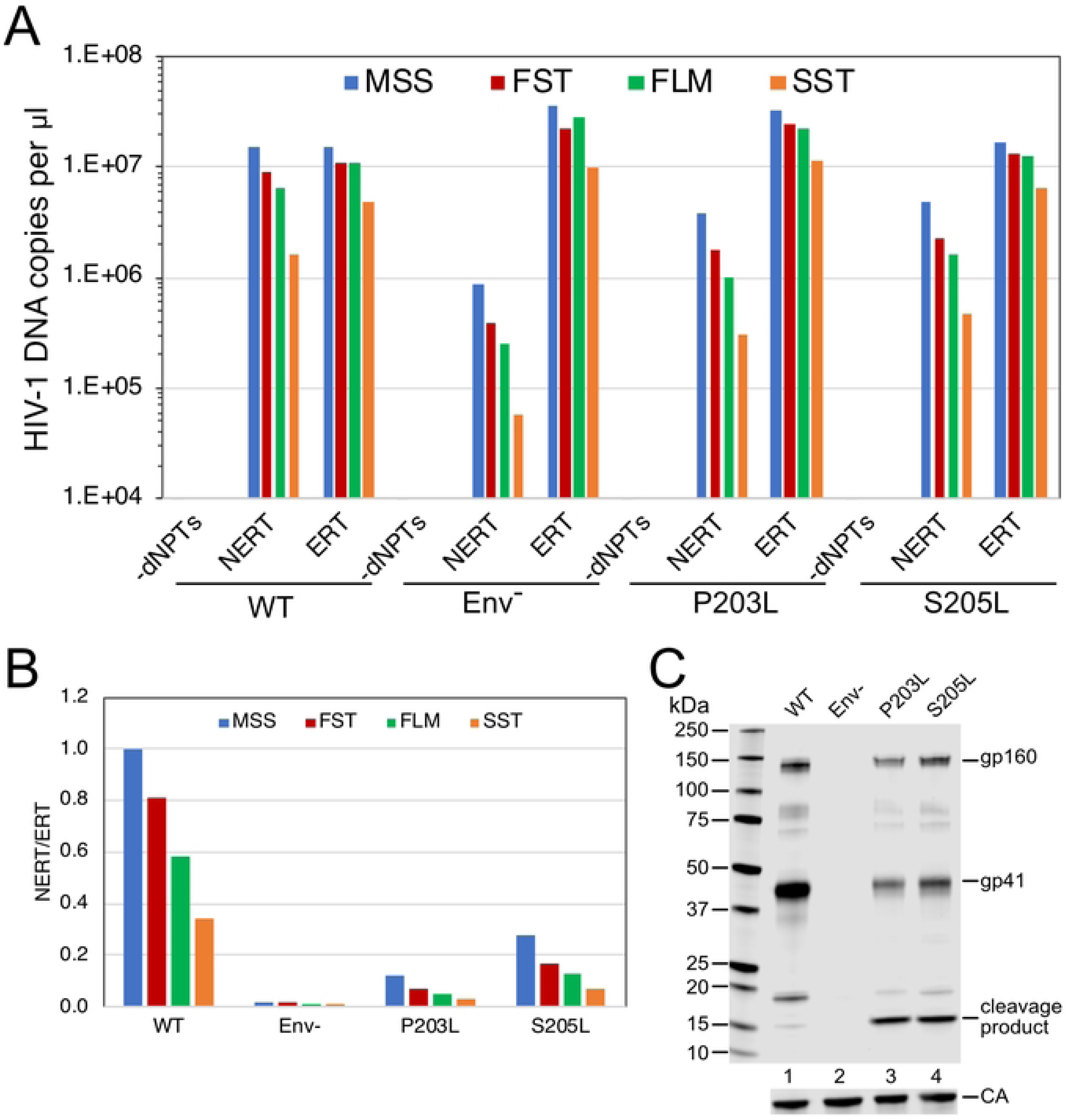
HIV-1 particles with cleavable gp41 CT proteins exhibit reduced NERT activity. A. Reactions were performed with the indicated wild type, Env-, and CT point mutant virions. Shown are the early (MSS and FST) and late (FLM and SST) products from the corresponding NERT and ERT reactions. B. Ratios of the DNA levels detected in the NERT and ERT reactions shown in A. C. Immunoblot analysis of the concentrated particles used in this experiment. Results shown are from one of two independent experiments.

### NERT activity does not require the mature viral capsid

In a previous study, we showed that PF74 binds specifically and stably to purified HIV-1 particles (15), indicating that the compound can cross the viral membrane. Therefore, the observed independence of NERT on IP6 and resistance to the capsid-targeting inhibitor PF74 (Fig 3) suggested that NERT does not require a stable viral capsid, unlike ERT reactions in permeabilized virions and purified cores. To further test whether formation of the mature HIV-1 capsid is required for NERT activity, we performed reactions with particles that were arrested in maturation via amino acid substitutions that block cleavage of various sites in the Gag polyprotein by the viral protease (PR). Because such mutations do not affect processing of the *pol* open reading frame by PR, both IN and RT are produced in normal quantities. We observed that mutations preventing Gag cleavage at the MA-CA and CA-SP1 junctions did not affect NERT activity but reduced ERT nearly to background levels (Fig 9A). By contrast, the MA/p6 mutant virus, containing cleavage-blocking substitutions at each of the Gag cleavage sites, exhibited moderately reduced NERT activity. Additionally, mutant particles with large in-frame deletions in the N-terminal domain of CA retained NERT activity but were markedly impaired in ERT reactions (Fig 9B). Collectively, these results show that efficient reverse transcription in intact HIV-1 particles does not require the mature HIV-1 capsid.

**Fig 9.**
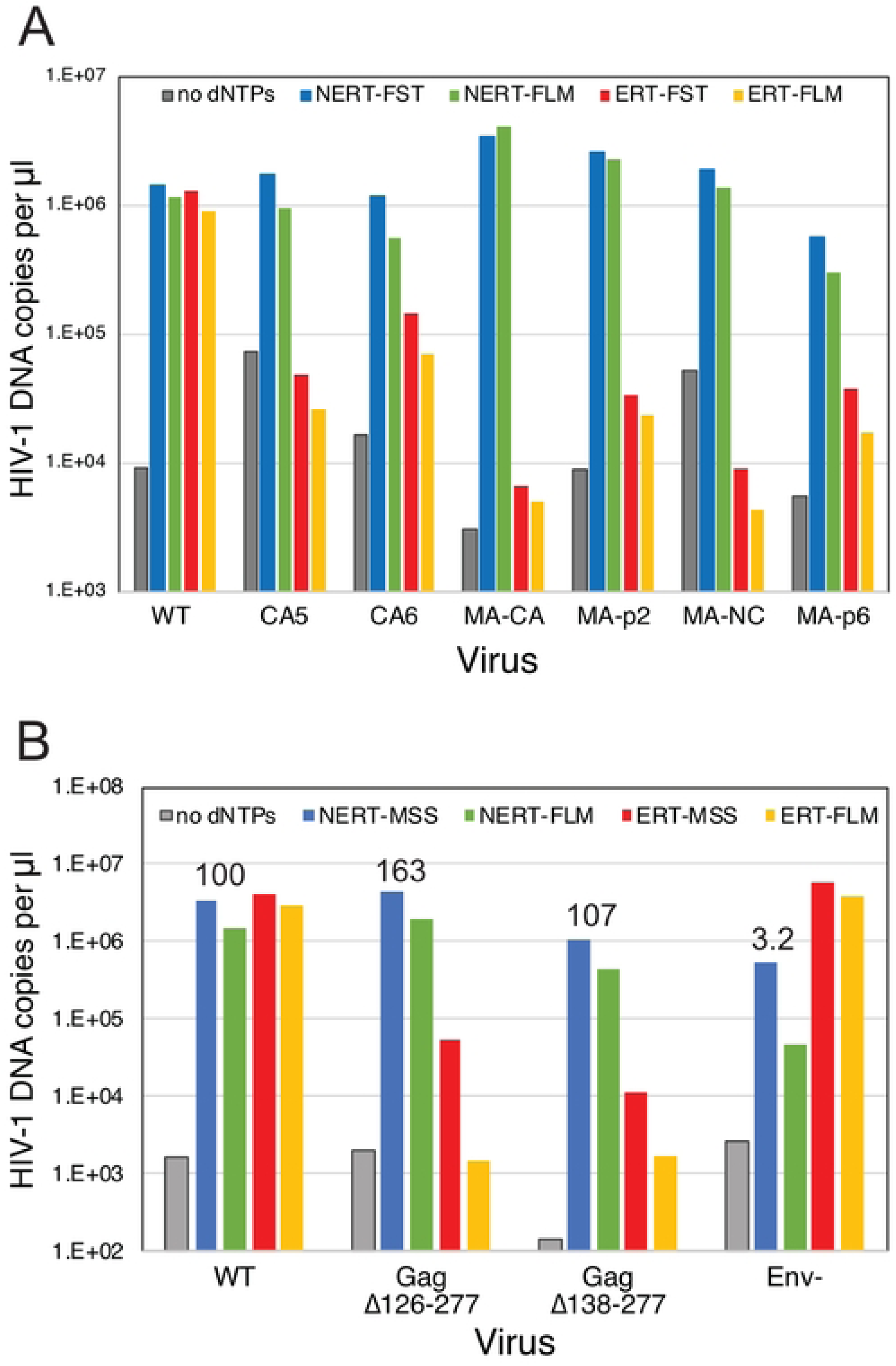
NERT activity occurs in particles lacking a mature capsid. A. NERT and ERT assays of the indicated wild type (NL4-3) and Gag cleavage mutants. CA5 particles contain uncleaved CA-SP1 protein; CA6 particles contain uncleaved CA-SP1-NC; MA-CA contains uncleaved MA-CA protein; MA-p2: uncleaved MA-CA-SP1; MA-NC: uncleaved MA-CA-SP1-NC; and MA-p6: uncleaved MA-CA-SP1-NC-SP1-p6. Shown are the early and late product DNA levels (FST and FLM, respectively). B. HIV-1 mutants bearing large deletions in CA are competent for NERT. Two mutants lacking nearly the entire N-terminal domain of CA were assayed for NERT and ERT with wild type and Env-particles. Numerical values shown represent the relative efficiency of ERT (FLM products normalized by exogenous RT activity levels in the viruses). The results shown are from one of two independent experiments.

## Discussion

In this study, we sought to define the mechanism by which the viral capsid promotes HIV-1 reverse transcription. Previous studies demonstrated a strong enhancement of ERT by IP6 and linked this effect to capsid stabilization by the metabolite. Here we provide two lines of evidence that IP6 stabilizes the closed form of the capsid and that this form is associated with efficient ERT. First, we showed that the IP6 enhancement of ERT is associated with acquisition of partial resistance to DNAse I. ERT levels were enhanced by increasing IP6 concentrations, and the reaction products were largely refractory to degradation by the nuclease. The DNase I-resistance of the IP6-stimulated ERT products suggested that the reaction takes place within a sealed container, but it did not exclude the additional possibility of a scaffolding role of the capsid.

The second line of evidence for the container model for capsid function was based on the fortuitous observation that the ERT reaction can occur within nonpermeabilized virions. Zhang and coworkers had reported in the mid 1990s that addition of dNPTs and polycations such as spermidine to HIV-1 particles resulted in detectable levels of DNA synthesis (31). They termed this reaction “NERT” for Natural Endogenous Reverse Transcription and proposed that this activity could promote sexual transmission of HIV-1. By contrast to the earlier study, our NERT reactions occurred efficiently in the absence of added polycations. Remarkably, unlike ERT, the NERT reaction did not require IP6, was resistant to PF74, and occurred in virions containing incompletely matured cores and in deletion mutants lacking the N-terminal domain of CA. We conclude that the NERT reaction is essentially independent of the viral capsid. Therefore, the intact viral membrane can serve as a container for the reverse transcription reaction in lieu of the mature capsid. The lack of a requirement for a mature capsid in the NERT reaction indicates that a scaffolding function of the capsid does not substantially contribute to efficient HIV-1 reverse transcription, at least *in vitro*.

If the HIV-1 capsid serves as a container for reverse transcription, what must be contained? One likely candidate is RT. Because the processivity of RT is relatively low, the local concentration of this enzyme in proximity to the viral genome may be critical. In preliminary studies, we have observed that addition of purified reverse transcriptase to ERT reactions lacking IP6 results in only a modest increase in late product synthesis. Based on the calculated volume of HIV-1 viral core (32), the reported proportion of Gag and Gag-Pol proteins in virions, and assuming that approximately one half of the RT resides within the core, we estimate that the RT concentration within an average core is roughly 0.2 mM—far higher than that which we added to the reactions. Nonetheless, the ability of purified RT to saturate reactions containing high concentrations of artificial substrates suggested that it is not limiting and that components of the viral core must be retained. One plausible candidate is the nucleocapsid protein which plays critical roles in the primer placement and strand transfer steps of reverse transcription (33). ERT reactions may be useful for studying the role of NC proteins in HIV-1 reverse transcription in the context of the viral core.

We also confirmed the previous observation by Zhang and coworkers that NERT is dependent on the HIV-1 Env complex and that truncation of the gp41 CT reduces NERT activity. The mechanism by which the viral Env protein renders virions permeable to dNTPs remains to be determined. Protease shaving of intact HIV-1 particles reduced NERT activity, and two HIV-1 mutants with full-length Env proteins bearing single amino acid substitutions in the gp41 CT resulting in its cleavage by the viral protease exhibited reduced NERT. A simple explanation for these results is that the Env protein complex itself, or a polypeptide derived from it (34), forms a pore in the viral membrane that permits passage of dNTPs and other small molecules. The gp41 CT contains membrane-interacting domains termed lentivirus lytic peptides that are capable of disrupting membranes when added as synthetic peptides (35), suggesting that these regions of the trimeric Env protein complex may form structures that permeabilize the viral membrane. Although we cannot formally exclude the possible involvement of a virion-incorporated host cell protein in NERT, the reduced activity observed in virions in which the CT is cleaved by the viral protease suggests otherwise. In any event, our results argue against nonspecific physical damage to the membrane that may occur during virion processing as a plausible explanation for virion permeability to dNTPs.

The ability of HIV-1 Env to render the viral membrane permeable to dNTPs suggests the possibility of a previously unrecognized role of the gp41 CT in HIV-1 replication and/or pathogenesis. Lentiviral Env transmembrane proteins are typified by long CTs, and while the gp41 CT is known to have important trafficking and regulatory functions, these are apparently served by shorter sequences in other retroviral genera (36). Membrane permeabilization by Env may prime virions for reverse transcription prior to cell entry, thereby increasing the fraction of particles that complete the process following membrane fusion with target cells. Additionally, membrane permeabilization by Env on the cell surface may promote the release of small molecules from infected cells, potentially promoting inflammation and/or priming of neighboring cells for infection. Finally, deposition of Env on the surface of cells via virion fusion with the plasma membrane could render target cells permeable to extracellular dNTPs, thus promoting reverse transcription in resting T cells that contain low levels of nucleotides.

The NERT reaction may also have practical utility. HIV-1 particles contain low levels of Env, and rapid and sensitive assays to determine the susceptibility of variants in patients to neutralizing antibodies are needed to guide selection of the appropriate therapeutic antibodies for individuals (37). However, detection of functional HIV-1 particles in patients may be confounded by the heterogeneity of Env levels on particles, which are typically low (38). The NERT reaction may provide a rapid and sensitive approach to enrich for genomes from HIV-1 particles bearing functional Env proteins in patient samples containing both defective and infectious HIV-1 particles.

## Materials and Methods

### Plasmids and viruses

Wild type HIV-1 particles were produced by transfection of the full-length pNL4-3 molecular clone (39) or its corresponding mutants. CT truncation mutants and CA partial deletion mutants (Δ126-277 and Δ138-277) were generated by PCR overlap cloning and modeled after mutants characterized in a previous study (40). Mutant plasmids were verified by Sanger sequencing of the PCR-generated regions. The Gag cleavage mutants were previously described (41, 42). pNL4-3-based chimeric clones encoding primary isolate Env proteins were provided by Dr. Paul Bieniasz (43). The P203L and S205L CT mutants (29) were the generous gift of Dr. Eric Freed (National Cancer Institute). Plasmids pHCMV-G (44) and pSV-A-MLV-env (45), encoding VSV-G and A-MLV Env proteins, were obtained from Dr. Jane Burns and Dr. Nathaniel Landau, respectively. Plasmid pBR239E encoding full-length SIVmac239 was provided by Dr. Toshiaki Kodama. The *env*-defective mutant was constructed by end-filling of the HindIII site at nucleotide 7078, thus creating a frameshift in *env* at codon 74.

Viruses were produced by transfection of 293T cells in 100 mm dishes using polyethyleneimine (PEI). DNA-PEI mixtures were added to cultures and incubated overnight. The culture medium was aspirated, monolayers gently rinsed with 5 ml of PBS, and 7 ml of fresh culture medium added. After 24h of further culture, the virus-containing supernatants were harvested, clarified by passage through 0.45 μM pore-size syringe filters, and frozen in 1 ml aliquots at -80°C.

Experiments involving treatment of virions with proteinase K were performed as follows. Proteinase K was added to concentrated DNase I-treated virions to a final concentration of 0.1 mg/ml, incubated for 1h at 37°C, and the virions pelleted through 20% sucrose to remove the protease. Virions were redissolved in their original volume and added to NERT reactions. Samples were also subjected to SDS-PAGE and immunoblotting.

For ERT and NERT reactions, aliquots of viruses were thawed in a 37°C water bath and MgCl_2_ and DNase I were added to 10 mM and 20 μg/ml concentrations, respectively. Following incubation at 37°C for 1h, the particles were pelleted through a 0.25 ml cushion of STE buffer (20 mM Tris-HCl pH 7.5, 150 mM NaCl, 1 mM EDTA) containing 20% sucrose by ultracentrifugation for 30 min at 45,000 rpm in a Beckman TLA-55 rotor. The supernatants were carefully removed by aspiration and the pellets resuspended in 50 μl of STE buffer.

### Endogenous reverse transcription reactions

Reactions were performed in 50 μl volumes containing 20 mM Tris-HCl pH 7.6, 2 mM MgCl_2_, 150 mM NaCl, 1 mM DTT, 1 mg/ml bovine serum albumin, 0.1 mM each dNTP, 0.1% (vol/vol) Triton X-100 (ERT) and the indicated concentrations of IP6 (TCI America). Reactions were initiated by the addition of virions and incubated at 37°C for either 4h or 14h. DNA was purified from the reactions using silica columns and assayed for early and late reverse transcripts by qPCR as previously described (20). The following compounds were tested: azidothymidine triphosphate (AZTTP, Jena Bioscience); efavirenz (EFV, HRP cat. no 4624); stavudine triphosphate (d4TTP, Jena Bioscience), PF-3450074 (PF74, MedChemExpress); aldrithiol (AT-2, Sigma).

### Immunoblotting

Pelleted virions were dissolved in SDS-PAGE sample buffer and subjected to electrophoresis on precast 4-20% gradient gels using Tris-MOPS running buffer (Genscript). Proteins were transferred to Protran nitrocellulose membranes (Perkin-Elmer) with a Genie electroblotter (Idea Scientific). Blots were blocked for 1h with 5% nonfat dry milk dissolved in PBS and probed with 1 μg/ml anti-gp41 monoclonal antibody (Chessie 8) produced in-house from the corresponding hybridoma. Proteins were detected with a IR680 dye-conjugated anti-mouse secondary antibody. Bands were revealed by scanning the blot with a LI-COR Odyssey imager and quantified with the instrument software. Following reprobing with an anti-CA monoclonal antibody (produced from hybridoma186-H12-5C), the relative ratios of gp41 to CA were calculated from the quantified band intensities.

## Acknowledgments

We thank the members of the Aiken lab for helpful suggestions during the course of this work. The following reagents were obtained through the NIH HIV Reagent Program, Division of AIDS, NIAID, NIH: Human Immunodeficiency Virus 1 (HIV-1), strain NL4-3 Infectious Molecular Clone (pNL4-3), ARP-114, contributed by Dr. M. Martin; anti-Human Immunodeficiency Virus 1 (HIV-1) gp120 Hybridoma (Chessie 8), ARP-526, contributed by Dr. George K. Lewis; anti-Human Immunodeficiency Virus 1 (HIV-1) p24 Hybridoma (183-H12-5C), ARP-1513, contributed by Dr. Bruce Chesebro and Dr. Hardy Chen. Supported by NIH grant R01 AI157843. AVR was supported by a CFAR Diversity, Equity, and Inclusion Pathway Initiative (CDEIPI) supplement to P30AI036219.

## References

1. Campbell EM, Hope TJ. 2015. HIV-1 capsid: the multifaceted key player in HIV-1 infection. Nat Rev Microbiol 13:471–83.

2. Zila V, Muller TG, Muller B, Krausslich HG. 2021. HIV-1 capsid is the key orchestrator of early viral replication. PLoS Pathog 17:e1010109.

3. Tailor MW, Chahine EB, Koren D, Sherman EM. 2023. Lenacapavir: A Novel Long-Acting Capsid Inhibitor for HIV. Ann Pharmacother doi:10.1177/10600280231171375:10600280231171375.

4. Forshey BM, von Schwedler U, Sundquist WI, Aiken C. 2002. Formation of a human immunodeficiency virus type 1 core of optimal stability is crucial for viral replication. J Virol 76:5667–77.

5. Tang S, Murakami T, Agresta BE, Campbell S, Freed EO, Levin JG. 2001. Human immunodeficiency virus type 1 N-terminal capsid mutants that exhibit aberrant core morphology and are blocked in initiation of reverse transcription in infected cells. J Virol 75:9357–66.

6. Yufenyuy EL, Aiken C. 2013. The NTD-CTD intersubunit interface plays a critical role in assembly and stabilization of the HIV-1 capsid. Retrovirology 10:29.

7. Reicin AS, Ohagen A, Yin L, Hoglund S, Goff SP. 1996. The role of Gag in human immunodeficiency virus type 1 virion morphogenesis and early steps of the viral life cycle. J Virol 70:8645–52.

8. Dorfman T, Bukovsky A, Ohagen A, Hoglund S, Gottlinger HG. 1994. Functional domains of the capsid protein of human immunodeficiency virus type 1. J Virol 68:8180–7.

9. Wang CT, Barklis E. 1993. Assembly, processing, and infectivity of human immunodeficiency virus type 1 gag mutants. J Virol 67:4264–4273.

10. Furuta RA, Shimano R, Ogasawara T, Inubushi R, Amano K, Akari H, Hatanaka M, Kawamura M, Adachi A. 1997. HIV-1 capsid mutants inhibit the replication of wild-type virus at both early and late infection phases. FEBS Lett 415:231–4.

11. Ganser-Pornillos BK, Pornillos O. 2019. Restriction of HIV-1 and other retroviruses by TRIM5. Nat Rev Microbiol 17:546–556.

12. Dismuke DJ, Aiken C. 2006. Evidence for a functional link between uncoating of the human immunodeficiency virus type 1 core and nuclear import of the viral preintegration complex. J Virol 80:3712–20.

13. Yang R, Shi J, Byeon IJ, Ahn J, Sheehan JH, Meiler J, Gronenborn AM, Aiken C. 2012. Second-site suppressors of HIV-1 capsid mutations: restoration of intracellular activities without correction of intrinsic capsid stability defects. Retrovirology 9:30.

14. Matreyek KA, Yucel SS, Li X, Engelman A. 2013. Nucleoporin NUP153 phenylalanine-glycine motifs engage a common binding pocket within the HIV-1 capsid protein to mediate lentiviral infectivity. PLoS Pathog 9:e1003693.

15. Shi J, Zhou J, Shah VB, Aiken C, Whitby K. 2011. Small-molecule inhibition of human immunodeficiency virus type 1 infection by virus capsid destabilization. J Virol 85:542–9.

16. Saito A, Ferhadian D, Sowd GA, Serrao E, Shi J, Halambage UD, Teng S, Soto J, Siddiqui MA, Engelman AN, Aiken C, Yamashita M. 2016. Roles of Capsid-Interacting Host Factors in Multimodal Inhibition of HIV-1 by PF74. J Virol 90:5808–5823.

17. Price AJ, Jacques DA, McEwan WA, Fletcher AJ, Essig S, Chin JW, Halambage UD, Aiken C, James LC. 2014. Host cofactors and pharmacologic ligands share an essential interface in HIV-1 capsid that is lost upon disassembly. PLoS Pathog 10:e1004459.

18. Balasubramaniam M, Zhou J, Addai A, Martinez P, Pandhare J, Aiken C, Dash C. 2019. PF74 Inhibits HIV-1 Integration by Altering the Composition of the Preintegration Complex. J Virol 93.

19. Christensen DE, Ganser-Pornillos BK, Johnson JS, Pornillos O, Sundquist WI. 2020. Reconstitution and visualization of HIV-1 capsid-dependent replication and integration in vitro. Science 370:eabc8420.

20. Jennings J, Shi J, Varadarajan J, Jamieson PJ, Aiken C. 2020. The Host Cell Metabolite Inositol Hexakisphosphate Promotes Efficient Endogenous HIV-1 Reverse Transcription by Stabilizing the Viral Capsid. mBio 11:e02820–20.

21. Sowd GA, Shi J, Fulmer A, Aiken C. 2023. HIV-1 capsid stability enables inositol phosphate-independent infection of target cells and promotes integration into genes. PLoS Pathog 19:e1011423.

22. Mallery DL, Faysal KMR, Kleinpeter A, Wilson MSC, Vaysburd M, Fletcher AJ, Novikova M, Bocking T, Freed EO, Saiardi A, James LC. 2019. Cellular IP6 Levels Limit HIV Production while Viruses that Cannot Efficiently Package IP6 Are Attenuated for Infection and Replication. Cell Rep 29:3983–3996 e4.

23. Sowd GA, Shi J, Aiken C. 2021. HIV-1 CA Inhibitors Are Antagonized by Inositol Phosphate Stabilization of the Viral Capsid in Cells. J Virol 95:e0144521.

24. Hu WS, Hughes SH. 2012. HIV-1 reverse transcription. Cold Spring Harb Perspect Med 2.

25. Jacques DA, McEwan WA, Hilditch L, Price AJ, Towers GJ, James LC. 2016. HIV-1 uses dynamic capsid pores to import nucleotides and fuel encapsidated DNA synthesis. Nature 536:349–53.

26. Mallery DL, Marquez CL, McEwan WA, Dickson CF, Jacques DA, Anandapadamanaban M, Bichel K, Towers GJ, Saiardi A, Bocking T, James LC. 2018. IP6 is an HIV pocket factor that prevents capsid collapse and promotes DNA synthesis. Elife 7:e35335.

27. Rossio JL, Esser MT, Suryanarayana K, Schneider DK, Bess JW, Jr., Vasquez GM, Wiltrout TA, Chertova E, Grimes MK, Sattentau Q, Arthur LO, Henderson LE, Lifson JD. 1998. Inactivation of human immunodeficiency virus type 1 infectivity with preservation of conformational and functional integrity of virion surface proteins. J Virol 72:7992–8001.

28. Zhang H, Dornadula G, Alur P, Laughlin MA, Pomerantz RJ. 1996. Amphipathic domains in the C terminus of the transmembrane protein (gp41) permeabilize HIV-1 virions: a molecular mechanism underlying natural endogenous reverse transcription. Proc Natl Acad Sci U S A 93:12519–24.

29. Waheed AA, Ablan SD, Roser JD, Sowder RC, Schaffner CP, Chertova E, Freed EO. 2007. HIV-1 escape from the entry-inhibiting effects of a cholesterol-binding compound via cleavage of gp41 by the viral protease. Proc Natl Acad Sci U S A 104:8467–71.

30. Waheed AA, Ablan SD, Mankowski MK, Cummins JE, Ptak RG, Schaffner CP, Freed EO. 2006. Inhibition of HIV-1 replication by amphotericin B methyl ester: selection for resistant variants. J Biol Chem 281:28699–711.

31. Zhang H, Dornadula G, Pomerantz RJ. 1996. Endogenous reverse transcription of human immunodeficiency virus type 1 in physiological microenviroments: an important stage for viral infection of nondividing cells. J Virol 70:2809–24.

32. Bryer AJ, Hadden-Perilla JA, Stone JE, Perilla JR. 2019. High-Performance Analysis of Biomolecular Containers to Measure Small-Molecule Transport, Transbilayer Lipid Diffusion, and Protein Cavities. J Chem Inf Model 59:4328–4338.

33. Rene B, Mauffret O, Fosse P. 2018. Retroviral nucleocapsid proteins and DNA strand transfers. Biochim Open 7:10–25.

34. Pfeiffer T, Ruppert T, Schaal H, Bosch V. 2013. Detection and initial characterization of protein entities consisting of the HIV glycoprotein cytoplasmic C-terminal domain alone. Virology 441:85–94.

35. Costin JM, Rausch JM, Garry RF, Wimley WC. 2007. Viroporin potential of the lentivirus lytic peptide (LLP) domains of the HIV-1 gp41 protein. Virol J 4:123.

36. Tedbury PR, Freed EO. 2015. The cytoplasmic tail of retroviral envelope glycoproteins. Prog Mol Biol Transl Sci 129:253–84.

37. Sneller MC, Blazkova J, Justement JS, Shi V, Kennedy BD, Gittens K, Tolstenko J, McCormack G, Whitehead EJ, Schneck RF, Proschan MA, Benko E, Kovacs C, Oguz C, Seaman MS, Caskey M, Nussenzweig MC, Fauci AS, Moir S, Chun TW. 2022. Combination anti-HIV antibodies provide sustained virological suppression. Nature 606:375–381.

38. Zhu P, Chertova E, Bess J, Jr., Lifson JD, Arthur LO, Liu J, Taylor KA, Roux KH. 2003. Electron tomography analysis of envelope glycoprotein trimers on HIV and simian immunodeficiency virus virions. Proc Natl Acad Sci U S A 100:15812–7.

39. Adachi A, Gendelman HE, Koenig S, Folks T, Willey R, Rabson A, Martin MA. 1986. Production of acquired immunodeficiency syndrome-associated retrovirus in human and nonhuman cells transfected with an infectious molecular clone. J Virol 59:284–91.

40. Borsetti A, Ohagen A, Gottlinger HG. 1998. The C-terminal half of the human immunodeficiency virus type 1 Gag precursor is sufficient for efficient particle assembly. J Virol 72:9313–7.

41. Wiegers K, Rutter G, Kottler H, Tessmer U, Hohenberg H, Krausslich H-G. 1998. Sequential steps in human immunodeficiency virus particle maturation revealed by alterations of individual Gag polyprotein cleavage sites. J Virol 72:2846–2854.

42. Wyma DJ, Jiang J, Shi J, Zhou J, Lineberger JE, Miller MD, Aiken C. 2004. Coupling of human immunodeficiency virus type 1 fusion to virion maturation: a novel role of the gp41 cytoplasmic tail. J Virol 78:3429–35.

43. Zhang YJ, Hatziioannou T, Zang T, Braaten D, Luban J, Goff SP, Bieniasz PD. 2002. Envelope-dependent, cyclophilin-independent effects of glycosaminoglycans on human immunodeficiency virus type 1 attachment and infection. J Virol 76:6332–43.

44. Yee JK, Friedmann T, Burns JC. 1994. Generation of high-titer pseudotyped retroviral with very broad host range. Methods Cell Biol 43:99–112.

45. Landau NR, Page KA, Littman DR. 1991. Pseudotyping with human T-cell leukemia virus type I broadens the human immunodeficiency virus host range. J Virol 65:162–169.

